# TissueMosaic enables cross-sample differential analysis of spatial transcriptomics datasets through self-supervised representation learning

**DOI:** 10.1101/2024.11.07.622479

**Authors:** Sandeep Kambhampati, Luca D’Alessio, Fedor Grab, Stephen Fleming, Fei Chen, Mehrtash Babadi

**Affiliations:** Bioinformatics and Integrative Genomics Program, Harvard Medical School, Boston, MA 02115, USA; Broad Institute of MIT and Harvard, Cambridge, MA 02142, USA; Data Sciences Platform, Broad Institute of MIT and Harvard, Cambridge, MA 02142, USA; Department of Stem Cell and Regenerative Biology, Harvard University, Cambridge, MA 02142, USA

## Abstract

Spatial transcriptomics allows for the measurement of gene expression within native tissue context, thereby improving our understanding of how cell states are modulated by their microenvironment. Despite technological advancements, computational methods to link cell states with their microenvironment and perform comparative analysis across different samples and conditions are still underdeveloped. To address this, we introduce TissueMosaic (*Tissue MOtif-based SpAtial Inference across Conditions*), a self-supervised convolutional neural network designed to discover and represent tissue architectural motifs from multi-sample spatial transcriptomic datasets (https://github.com/broadinstitute/TissueMosaic). TissueMosaic effectively maps structurally similar tissue motifs close together in a learned latent space. TissueMosaic further links these motifs to gene expression, enabling the study of how changes in tissue structure impact function. TissueMosaic increases the signal-to-noise ratio of differential expression analysis through a motif enrichment strategy, resulting in more reliable detection of genes that covary with tissue structure. Here, we demonstrate TissueMosaic on high resolution spatial transcriptomics datasets across tissues, learning representations that outperform neighborhood cell-type composition baselines and existing methods on downstream tasks. We highlight genes and pathways in these tissues that are associated with changes in tissue structure across external conditions. These findings underscore the potential of self-supervised learning to significantly advance spatial transcriptomics research.

## Introduction

Spatial transcriptomics enables the measurement of gene expression in the native context of tissues. It thus serves as a powerful experimental platform to understand how cell state is modulated through cell non-autonomous effects. Despite the rapid progress of spatial transcriptomics (ST) technologies in recent years, effective computational approaches to infer the relationship between the cell state and its microenvironment are still lagging behind, and it remains difficult to learn the functional organization of tissues from ST data. A conceptual barrier is that analyzing how gene expression is coupled to changes in tissue structure first requires the notion of a basic “unit” of tissue structure. This unit would allow the comparison of samples across multiple conditions by serving the function of a shared representation space. This would be analogous to how the basis of “genes” allows unambiguous quantification and comparison of bulk transcriptomics data across multiple conditions. We term these units *tissue motifs* and aim to discover these motifs in an unbiased way.

One logical way to represent these units would be with handcrafted features, such as the cell-type compositions of the local neighborhoods around each cell. However, handcrafted features are often biased and fail to fully represent the true sources of tissue structure variability. For example, the local neighborhood cell-type composition does not represent the geometric relationships between cells and cannot capture features like disorder or bunching. A common alternative approach, weakly-supervised representation learning, allows learning features that are best suited for a particular downstream task. Unfortunately, it is difficult and laborious to acquire labeled annotations for tissue structure. Furthermore, tissues are organized at multiple hierarchies, complicating such annotation efforts.

In many fields, such as computer vision, self-supervised learning (SSL) provided a paradigm shift by learning features that are data-driven and capture the complex underlying grammar of the raw data. SSL is a framework that leverages inherent features of the data to formulate a proxy supervised task. To solve this supervised task, the model is encouraged to extract information-rich and robust representations of the data. If the self-supervised objective is designed well,^1–3^ these representations can also generalize to multiple other downstream tasks. These features generally outperform handcrafted and weakly-supervised features on multiple downstream tasks^4^. Thus, to learn representations of tissue architecture in the absence of labeled histological motifs, we turn to self-supervised learning.

Here, we introduce TissueMosaic (*Tissue MOtif-based SpAtial Inference across Conditions*), a method that leverages self-supervised learning to learn unified representations of tissue architectural motifs from multi-sample spatial transcriptomic datasets. TissueMosaic provides a data-driven basis to quantify the relatedness and dissimilarity of local tissue motifs across different samples. It also provides a framework to connect these learned motifs back to gene expression. Through this method, we are able to specifically study how changes in tissue structure across different conditions impact tissue function. Critically, this approach allows us to identify and prioritize *condition-specific (such as disease-enriched)* tissue motifs, a task we call “conditional motif enrichment”. Conditional motif enrichment increases our power to perform spatial differential expression despite the spatial heterogeneity inherent to tissues.

## Results

### TissueMosaic learns representations of spatial transcriptomics data via self-supervised learning

Our self-supervised learning workflow is designed for learning representations of “tissue structure” from spot-based spatial transcriptomics data (such as Slide-seqV2^5^, Stereo-seq^6^, and Visium^7^) or cell-based spatial transcriptomics data (such as Slide-tags^8^). In these data, we refer to individual “spots” as gene expression profiles sampled from a given spatial location. We define tissue structure as sufficient statistics of the geometric relationship between “molecularly defined cell types” in space. Thus, we first obtain a lossy encoding of the raw gene expression data that reduces transcriptomic profiles to broader cell type or cell state labels. This low dimensional encoding could be the first few principal components of the measured gene expression, for example, or the cell-type proportions or cell-types inferred at each spot.^9^

Importantly, these broader labels are deliberately encoded at a coarse resolution relative to the underlying transcriptomic variability. This first has the advantage of mitigating technical batch effects, allowing us to learn robust representations that will generalize across samples. Second, this encoding results in spatial representations that do not contain information related to cell state variability within the broad cell type labels. We are thus able to subsequently use the spatial representations to study the covariation between the more detailed cell states and local tissue structure.

We then rasterize the coarse spot encodings and their spatial coordinates into a multi-channel image, with one channel per dimension of the spot encoding (e.g. inferred proportions of each cell-type, with typically less than 20 dimensions **Methods**). We refer to this processed data as “cell-type images”, which capture the spatial relationships critical to defining tissue structure. Cell-type images also enable easy integration with additional imaging modalities, such as Hematoxylin and eosin (H&E) staining or fluorescent imaging, that represent a complementary measurement of tissue structure.

Furthermore, cell-type images allow us to leverage abundant existing research on self-supervised learning frameworks for computer vision. In these frameworks, a common pretext task is designed around mapping similar data together in a latent space^10^. These similar instances are usually created by applying random augmentations to input images. The augmentations are carefully designed to retain the desired identity of each image while at the same time desensitizing the features to irrelevant information. The model is then trained to produce representations that are similar for each “positive pair” of augmented images. Various strategies are implemented to prevent collapse to a degenerate solution, such as using negative pairs^2^, redundancy reduction^11^, or sharpening and centering^12^.

We apply a similar strategy to train our model, randomly extracting patches of fixed size from the cell-type images as our model input. These patches define the unit of tissue structure from which tissue structure is compared across samples. The size of the patches is a hyperparameter of the model that is either set based on prior knowledge of the relevant length scale of a given tissue’s structure or by learning this length scale (**Methods**). We apply random augmentations to these patches and pass them as input to two different branches of the model (**Fig. 1A**). These augmentations include random flipping, rotation, dropout of spots, multiplication by random intensity, random rotation, random horizontal and vertical flipping, random resized crops, straight cuts to occlude parts of the image, and dropout of channels (cell-types) (**Fig. S1A, Methods**).

**Figure 1.**
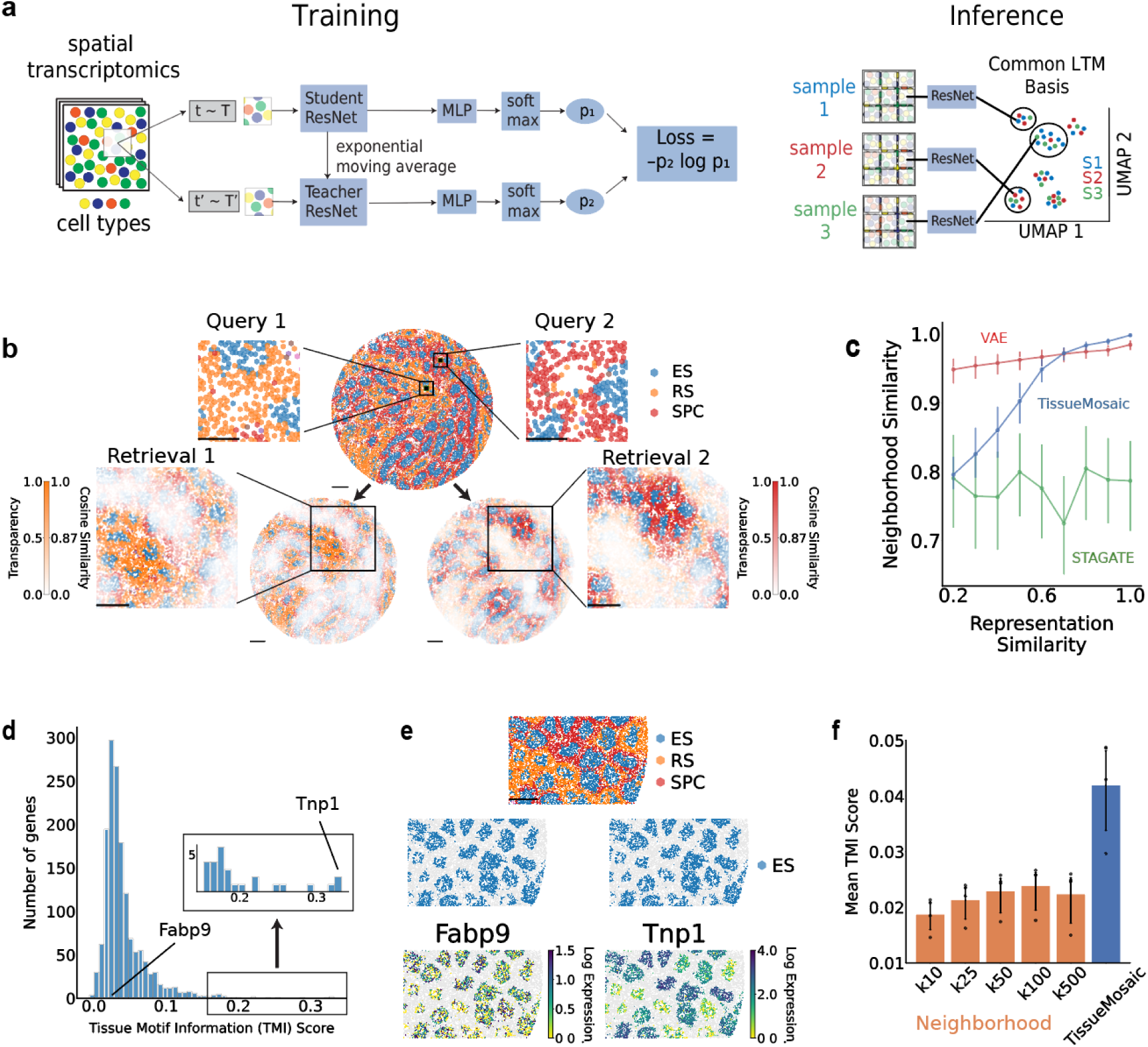
**a**. (Left) TissueMosaic trains a Residual Convolutional Neural Network (ResNet) via a self-supervised learning framework (DINO) to learn representations of spatial transcriptomics data cast as images. (Right) TissueMosaic learns a common representation space to summarize local tissue motifs (LTM) within a spatial transcriptomics dataset. **b.** Motif query on TissueMosaic representations of a Slide-seqV2 mouse testis dataset reveals tubules that are either rich in round spermatids or spermatocytes. For retrieval (bottom), exponentiated cosine similarity to the query patch is plotted as an alpha transparency mask (Methods). **c.** Performance of TissueMosaic, a Variational Autoencoder (VAE) baseline, and a previous representation learning method, STAGATE, across all testis samples on the motif query task, evaluated by correlation of cosine representation similarity with cosine neighborhood similarity. Median cosine neighborhood similarity in 0.1 intervals of cosine representation similarity is plotted. Error bars indicate 20% percentile intervals of the data. VAE *r* = 0.2696, TissueMosaic *r* = 0.4435, STAGATE *r* = –0.0062, across all data (Methods). **d.** Distribution of tissue motif information (TMI) scores of highly expressed genes in elongated spermatid cells. Inset shows genes with TMI greater than 0.15. **e.** Left, example of genes with low (left) and high (right) tissue motif information (TMI) scores, 0.029 and 0.31 respectively, in Elongated Spermatid (ES) cells in sample region of Slide-seqV2 mouse testis dataset. **f.** Performance of gene expression regression with neighborhood compositions across multiple neighborhood sizes and TissueMosaic representations, evaluated by mean TMI score. Error bars indicate 95% confidence interval (determined by bootstrapping) of the mean Tissue Motif Information (TMI) score from spatially partitioned test folds (N=4, Methods). Scale bars: **b)** query insets, 100 µm; all other scale bars 300 µm, **e)** 100 µm

After the model is trained, during inference time, we sample patches from our spatial transcriptomics samples, pass them through the model, and obtain a set of representations that encapsulate the local tissue organization within each sample (**Fig. 1A**). Our method, TissueMosaic, learns representations that enable diverse downstream tasks such as ***motif query***, ***spatial clustering***, and identification of ***tissue motif informed genes*** (**Fig. S1B**).

We evaluated three different self-supervised loss functions originally designed for learning representations of natural images: SimCLR^2^, Barlow Twins^11^, and DINO^12^ and found that on our cell-type images, DINO produces representations that are most informative of underlying tissue features and have the best performance on downstream tasks (**Fig. S2A, Fig. S4C**). We hypothesize that this is due to the local/global patch sampling augmentation that is unique to the DINO approach and encourages the model to learn local-to-global correspondences in the tissue architecture. A simple ablation study removing the local/global sampling strategy resulted in worsened quality of the learned representations (**Fig. S2B)**, highlighting the importance of learning local-to-global relationships towards understanding tissue organization.

### TissueMosaic representations are biologically meaningful with respect to underlying tissue structure

We first explored the representations of tissue motifs learned by TissueMosaic. In meaningful representations, we expect tissue patches containing visually similar motifs to lie close to each other in the learned latent space. Thus, we first examine the learned representations through the downstream tasks of ***motif query,*** i.e. retrieving structurally similar tissue patches to a query patch, and ***spatial clustering,*** i.e. grouping structurally similar tissue patches together.

We demonstrate these downstream tasks on two Slide-seqV2 datasets: (1) a mouse testis dataset, consisting of 3 wild-type samples and 3 ob/ob samples, representing a genetic mouse model of diabetes, with 187,422 total spots and (2) a mouse thymus dataset^13^, with a time course of 47 samples spanning Day 0 to Week 90 of mouse lifespan, with 1,255,473 total spots. We selected these datasets due to the presence of well defined architecture in these tissues that is closely related to biological function. The basic architectural unit of testis tissue is the seminiferous tubule, which depending on the developmental stage of the tubule, has differential cell type composition^14^. The maturity of germ cells is associated with their radial position within each tubule^14^. The thymus is separated into two main anatomical regions: the cortex and the medulla^15^. The spatial structure of the cortex versus medulla is important for the maturation of T cells in the thymus, and changes in this structure are linked to declining thymus function with age.^16^

We first applied TissueMosaic to the mouse testis dataset to use the learned representations to ***“query” spatial motifs***, i.e. find structurally similar tissue patches in the dataset given a reference tissue patch. In the testis data, there are two main tissue motifs: tubules rich in round spermatids (RS) and tubules rich in spermatocytes (SPC). Querying a sample with either of these motifs highlights the other portions of the sample that contain this motif (**Fig. 1B**).

Intriguingly, despite having no input information regarding tubule segmentation, the model is able to learn the boundaries between individual tubules (**Fig. 1B**). Within the latent space of the representations learned by the model, the proportion of round spermatids vs spermatocytes in each patch captures the main axis of variation (**Fig. S2A**). We quantify the model’s performance on the motif query task by computing whether query and retrieved patches that have similar TissueMosaic representations also have similar cell-type compositions, since cell-types drive the major motifs in this dataset (**Fig. 1C**). We observe that TissueMosaic has a stronger correlation between model representation similarity and neighborhood similarity (*r* = *0.4435*) compared to either our own variational autoencoder (VAE) baseline (*r = 0.2696*) or a previous autoencoding-based method, STAGATE^17^ (*r = –0.0062*). We note that in contrast to STAGATE, TissueMosaic is able to leverage information across all samples in the dataset, likely contributing to better performance.

We next applied TissueMosaic to the mouse thymus dataset to demonstrate that the learned representations enable ***spatial clustering***. On this dataset, we cluster the learned representations to separate the cortex and medulla (**Fig. S3A**). The latent space learned by the model shows strong separation based on the proportion of cortex beads and medullary beads (determined by manual annotation, **Methods**) within each patch, even though the cortex/medullary annotation is not used as input to the model (**Fig. S3B**). Accordingly, clustering the TissueMosaic representations leads to high agreement with manual annotations of the cortex and medulla (Methods). This is likely due to the model picking up on the arrangement of different cell-type compositions within each anatomical region. To quantify clustering performance, we compute the mutual information between the clustering labels and manual annotations as an evaluation metric. We achieve stronger concordance with the manual annotation compared to clustering based on the cell-type composition of each patch alone (“Neighborhood”). We are also able to outperform existing spatial domain detection methods such as SpaGCN^18^ and STAGATE^17^ on this dataset (**Fig. S3C**).

### TissueMosaic connects gene expression to tissue microenvironments

After establishing that TissueMosaic learns meaningful representations of tissue structure, we next aimed to leverage the learned representations to study ***how tissue structure is linked to gene expression***. The spatial context of a cell can impact its state, and this effect can be mediated by specific genes. We reasoned that the TissueMosaic representations could be used to identify such genes.

To do so, we designed a Poisson Generalized Linear Model (GLM) to analyze how the spatial milieu around a particular cell modulates its gene expression (**Fig. S4A, Methods**). We model the measured gene expression at a given spot as a Poisson distribution, the rate parameter of which is a function of the number of UMIs present at that spot (N_n_), the cell-type proportions at that spot, and a choice of spatial covariates that capture the local tissue structure surrounding that spot (**Fig. S4A**). This could be, for example, the neighborhood cell type composition or the TissueMosaic representation of a patch centered at that spot. We train a separate model per cell-type, assigning spots to cell-types based on the majority cell-type present at each spot. We compare the performance of this spatial GLM to a baseline GLM that only uses the number of UMIs and cell-type proportion information (i.e. without spatial covariates). We then determine which genes gain the greatest predictive power by including information on tissue motifs. We term such genes “tissue motif informed (TMI) genes”. We define the tissue motif information score for a given gene *g* as:

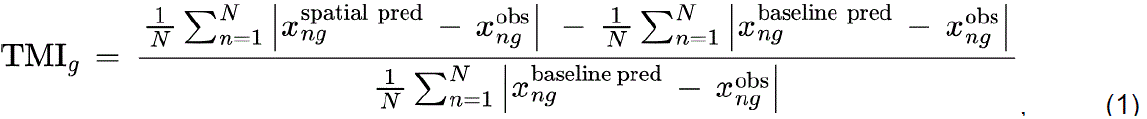

where *x* is the count of gene *g* at spot *n*,“obs” refers to observed (i.e. measured) counts, while “pred” refers to the predicted counts from the spatial or baseline regression model. We emphasize that because TMI genes are calculated in a cell-type specific manner and adjusted to a baseline cell-type GLM, they reflect spatial information *beyond* what cell type information alone could explain. Thus, TMI genes would not include marker genes of a given cell-type, but rather genes that have spatial variance *within* a cell-type due to the surrounding tissue structure.

Among the three major cell types in the testis data, elongated spermatid (ES) cells exhibit the highest mean tissue motif information score across highly expressed genes (**Fig. S5A**). The distribution of TMI scores in the testis for genes in ES cells (**Fig. 1D**) is skewed right, with most genes exhibiting low TMI scores but a minority with large TMI scores. This aligns with the expectation that only a subset of genes within a cell type will exhibit tissue structure informedness. Genes with high TMI scores, such as *Prm1* in ES cells, exhibit a highly structured spatial distribution of gene expression **(Fig. 1E)**. Crucially, the expression is also highly correlated with the tissue motif (Round Spermatid rich tubule or Spermatocyte rich tubule) in which the ES cell is located (**Fig. 1E, Fig. S5B)**. We highlight the distinction between spatially informed and spatially patterned gene expression. Although the spatial expression does not adhere to a simple gradient or pattern that can be easily parameterized, our framework still identifies the gene as spatially informative. In contrast, *Fabp9*, which has a tissue motif information score near the median of the distribution, shows a much more random spatial expression in ES cells **(Fig. 1E)**.

The mean tissue motif information score of highly expressed genes in ES cells using TissueMosaic representations far exceeds that of a baseline based on the cell-type compositions of the *k* nearest neighbors across multiple neighborhood sizes (“Neighborhood”, **Fig. 1F**). This indicates that information beyond the local neighborhood cell-type composition, including the geometric arrangement of neighboring cell types, is important for modeling the covariation of gene expression of a given cell and its microenvironment. We note similar trends for other key cell types in the testis (**Fig. S5A**).

In summary, TissueMosaic learns meaningful representations of spatial transcriptomics data that can be used to interrogate substructures of varying shapes and sizes within spatial transcriptomics data without any *a priori* knowledge about these structures. Furthermore, TissueMosaic links these representations back to gene expression to identify tissue motif informed genes.

### TissueMosaic enables tissue motif enrichment and differential expression analysis between enriched motifs

Beyond linking tissue motifs to genes informed by these motifs, we next aimed to use our framework to study how changes in tissue motifs due to external conditions co-vary with gene expression. Spatial transcriptomic datasets often come with additional metadata regarding experimental conditions, such as mutant vs. wild type (as in the mouse testis dataset), age (as in the mouse thymus dataset), etc. It is desirable to identify tissue motifs that serve as archetypes of each categorical condition or that covary with continuous conditions. One can then restrict spatial gene expression analysis to such archetypes as a strategy to focus on tissue regions of interest and to increase the statistical power in differential testing. We refer to this strategy as “conditional motif enrichment,” or simply motif enrichment.

To perform motif enrichment, we train a Ridge classifier or regressor to predict the sample condition label of individual cells from the learned TissueMosaic representations (**Fig. 2A, Methods**). We then use the trained classifier or regressor to predict the status of held out cells from each sample following a spatially partitioned train/test split (**Fig. S4B, Methods**), thereby distilling the sample into regions enriched for conditions present in the dataset. Our motif enrichment framework is related to previous work on weakly supervised methods, such as multiple instance learning^19^, but differs from such work, since the representations are learned directly from the data rather than external labels^20^.

**Figure 2.**
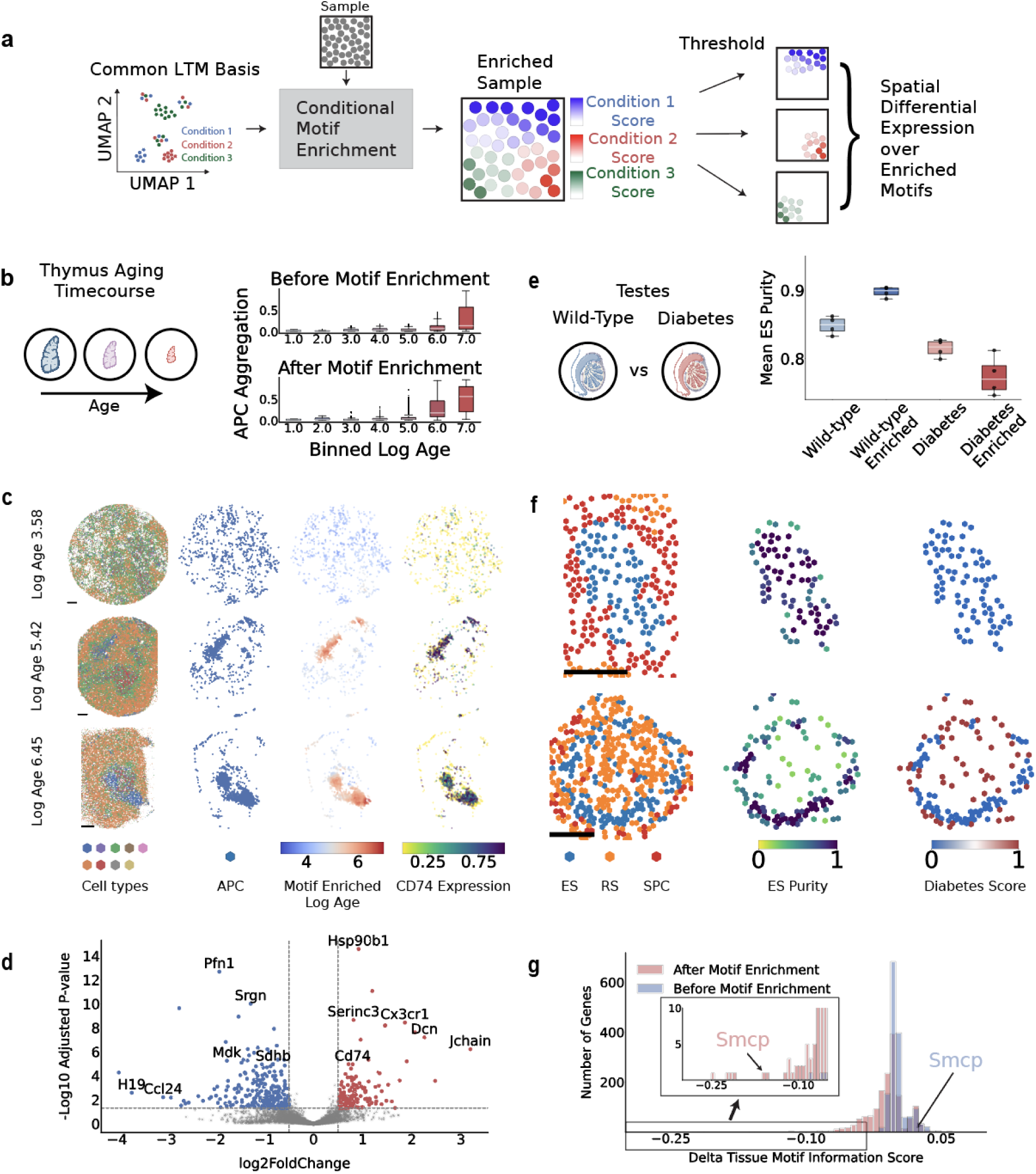
**a**. Conditional motif enrichment with TissueMosaic. Sample labels of conditions for input tissue motifs are used to train a classifier/regressor that scores held-out motifs on their enrichment for a given condition. A sample from a specific condition may contain motifs enriched for multiple conditions. **b.** Box plots of Antigen presenting cell (APC) aggregation scores, defined as proportion of APCs in k100 neighborhood, before and after motif enrichment. Log Age calculated as log2 of biological age in days. Center = median, Box = IQR, Whiskers = farthest data point within 1.5IQR from box. Data beyond 1.5IQR are plotted individually. Before motif enrichment, N = 247, 280, 699, 4077, 2042, 3546, 4268 cells per bin respectively. After motif enrichment, N = 13, 34, 109, 2088, 8713, 3279, 928 cells per bin respectively. **c.** Three thymus samples of different biological ages. Spots are colored by cell types (first and second columns), TissueMosaic trained regressor prediction of antigen presenting cell age (third column), and CD74 expression, normalized by library size and log transformed (fourth column). Spots in third and fourth columns are plotted with transparency of 0.5. Spots in the fourth column with CD74 expression > 0 are plotted with a higher z-order. **d.** Volcano plot of differential expression results between cells in motifs with motif enriched log age >= 5.0 vs cells in motifs with motif enriched log age < 5.0. Genes with adjusted p-value less than 0.05 and log fold change greater (lower) than 0.5 are highlighted in red (blue). **e.** Box plots of mean elongated spermatid purity per test-fold (N = 4), before and after motif enrichment. Center = median, Box = IQR, Whiskers = farthest data point within 1.5IQR from box. Data beyond 1.5IQR are plotted individually. **f.** Examples of organized (top) and disorganized (bottom) tubules. Left, cell-types localized in space, Middle, Elongated spermatid cells colored by ES purity (defined as proportion of elongated spermatid cells in k Nearest neighborhood). Right, TissueMosaic trained classifier prediction of elongated spermatid status. **g.** Distribution of differential tissue motif information scores ΔTMI (Methods) of highly expressed genes between wild-type motif enriched and diabetic motif enriched elongated spermatid cells. Inset shows genes with ΔTMI < –0.05. Scale bars: **c)** 300 µm, **f)** 100 µm

Conditional motif enrichment on a spatial transcriptomics dataset enables differential gene expression analysis between tissue motifs associated with external conditions. We demonstrate this in the thymus dataset, which is a time course of thymic involution, a gradual shrinking of the thymus with age,spanning Day 0 to Week 90. One feature of involution in the thymus is the spatial aggregation of antigen presenting cells (APCs) (**Fig. 2B**). However, only a subset of APCs are aggregated, even at later time points (**Fig. 2B**). To increase the biological signal of differential gene expression associated with APC aggregation, we first predict the biological age of APCs in each sample based on their motif enrichment scores.

In the thymus, motif enrichment distinguishes involuted vs non-involuted APCs based on their level of aggregation (**Fig. 2B**) and highlights the aggregated portions of the tissue as having high age (**Fig. 2C**). This in turn enables differential gene expression analysis specifically between the aged and non-aged cells within the tissues (**Fig. 2D)**. One gene highlighted by this analysis is *CD74*, whose overexpression is associated with B-cell malignancies^21^ (**Fig. 2C,D**). Gene set enrichment analysis (GSEA)^22^ of all significant DE genes, ranked by log-fold change, highlights various immune pathways, such as B cell activation (*FDR q-val = 0.159756*) and activation of immune response (*FDR q-val = 0.165778*) as upregulated in aged APCs (**Table S1**). This suggests that immune activation of APCs is associated with aggregation during thymus involution.

### Tissue motif enrichment detects greater effect size for genes with changes in TMI in condition-enriched motifs

To demonstrate the effectiveness of motif enrichment for differential analysis, we perform the following experiment in the mouse testis dataset. First, we identify genes whose TMI score changes between samples labeled for condition 1 vs condition 2. These are genes whose mutual information with tissue motifs covaries with external condition (beyond simply magnitude of expression). We then compare this analysis to the differential TMI score between enriched condition 1 vs condition 2 tissue motifs. We define the differential TMI score as:

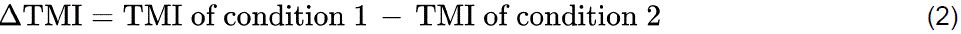

A gene with a large magnitude ΔTMI therefore has a substantial change in tissue motif information between condition 1 motifs and condition 2 motifs.

The mouse testis dataset consists of samples that are wild-type or from a mouse model of diabetes (ob/ob). In the ob/ob mice, a higher proportion of tubules exhibit spatial disorganization, indicated by a mixing of elongated spermatids (ES) cells with other cell types (we quantify this with an ES neighborhood purity score) (**Fig. 2E**). However, both organized and disorganized tubules are present in the wild-type and diabetic samples, though, in different proportions. Motif enrichment with the TissueMosaic representations increases the difference in ES purity scores between the enriched healthy vs enriched diseased cells compared to sample label alone (**Fig. 2E**). Notably, motif enrichment identifies organized vs disorganized ES cells at a sub-tubular level (**Fig. 2F**). To achieve this, we learn representations at a smaller patch size of 42 x 42 microns compared to the previous results on the testis data (patch size of 250 x 250 microns), since the representations at the larger patch size are unable to discern the sub-tubular disorganization (**Fig. S6A,B**).

We then train two independent GLMs on the healthy or diseased cells and compute the ΔTMI of genes between the healthy and diseased contexts. The ΔTMI between wild-type (condition 1) vs diabetic (condition 2) motifs in the testis forms a bi-modal distribution, with some genes better predicted in the wild-type context (Δ*TMI* > 0), but the majority of genes better predicted in the diabetic context (Δ*TMI* < 0) (**Fig. 2G**). We hypothesize that this is due to genes in the disorganized tubules exhibiting higher spatial variance, thus sharing more information with the TissueMosaic representations. These genes include *Smcp*, which was previously reported to be linked to the disorganization of tubules in diabetic testis.^23^ Intriguingly, *Smcp* only exhibits a large ΔTMI after motif enrichment (Δ*TMI = –0.00246*8 before motif enrichment, Δ*TMI = –0.1521* after motif enrichment). For other genes, such as *Tnp1*, *Tnp2*, *Prm1* and *Prm2*, motif enrichment increases the observed effect size in ΔTMI. (Before/after motif enrichment – Tnp1*: –0.06012 / –0.2127,* Tnp2*: –0.07960 / –0.2508,* Prm1*: –0.05349 / –0.2149,* Prm2*: –0.03483 / –0.2241*) Gene set enrichment analysis (GSEA) on ΔTMI after motif enrichment reveals that key processes in ES cells exhibit spatial dysregulation in diabetes, including processes related to DNA packaging complex (*FDR q-val = 0.01498*), Germ cell nucleus (*FDR q-val = 0.01902*), and Sperm motility (*FDR q-val = 0.09032*) (**Table S2**). These findings are consistent with previous studies suggesting a loss of ES identity in ob/ob testes,^23,24^ but further elucidates the spatial component of changes in ES gene expression in diabetes. This analysis demonstrates that motif enrichment can sufficiently increase signal-to-noise ratio to reliably detect genes that mediate the impact of tissue structure on cell state.

## Discussion

We introduce TissueMosaic, a method that learns unified representations for tissue architectural motifs across multi-sample spatial transcriptomic datasets via self-supervision. TissueMosaic links these representations back to gene expression and reveals how changes in tissue organization are coupled to alterations of transcriptional programs. TissueMosaic allows users to interrogate tissue structure and its relationship to gene expression, perform tissue motif enrichment, and conduct cross-sample spatial differential expression. Here, we demonstrate the method on two spatial transcriptomics datasets of mouse testis and thymus, illustrating that self-supervised learning (SSL) provides a powerful framework for learning representations of spatial transcriptomics data.

In our work, we specifically treat spatial transcriptomics data as images, a domain in which there is rich literature of SSL methods built on the backbone of convolutional neural networks (CNNs). Previous work has applied SSL to spatial transcriptomics using graph-based architectures.^25^ We reasoned that representing ST data as images provides a more expressive encoding of spatial relationships compared to graphs. The latter only captures local connectivity information but cannot capture higher order spatial features, such as disorder. Another advantage of treating tissue structure as images is the ease of multi-modal input to the model with other imaging modalities, such as fluorescent imaging or H&E staining. These images could simply be concatenated as additional channels in the input. It would be intriguing to leverage multi-modal input with TissueMosaic to learn richer representations of tissue structure beyond cell-type arrangements in space.

We recognize that image-based analysis of ST data has its own set of drawbacks. In particular, an image is an inefficient encoding of spatial transcriptomics data due to the typical sparsity of such data. Instead, using point cloud methods to encode and model the data would allow for more compute-efficient point cloud convolutions.^26,27^ Applying SSL to point clouds would be a natural evolution of TissueMosaic that would allow for analysis of larger fields of view by virtue of greater efficiency.

A limitation of our approach is the need to specify a fixed patch size as a model parameter. The patch sizes chosen for the datasets analyzed in this study reflects prior knowledge of the relevant length scales in a tissue and task specific manner. When such prior knowledge is not available, one can train TissueMosaic models with multiple patch sizes (as independent model training runs), perform the gene regression task with the resulting representations, and choose a patch size that explains the greatest tissue motif information. Future work could leverage the hierarchical nature of tissue structure to explicitly learn multi-scale representations of spatial transcriptomics data via an appropriately designed self-supervision objective. As the corpus of spatial transcriptomic data scales, such models, including TissueMosaic, will be critically needed for meaningful representations and analyses of tissues.

## Code Availability

TissueMosaic is implemented as an open-source python package, with code available at https://github.com/broadinstitute/TissueMosaic. We have provided a complementary codebase, https://github.com/broadinstitute/TissueMosaic_Manuscript to reproduce all of the figures in the manuscript. Comprehensive documentation and a quick-start tutorial demonstrating how to run the package on the Slide-seqV2 mouse testis dataset is available at https://tissuemosaic.readthedocs.io/.

## Data Availability

Pre-processed Slide-seqV2 data of the mouse testis is available at https://drive.google.com/file/d/1buvC7H8-EsyLrniMCre7Y7PZLu8zw8h2/view. Raw and preprocessed Slide-seqV2 data of the mouse thymus will be made publicly available upon publication of the corresponding manuscript.

## Acknowledgements

We thank Sophia Liu and Yerdos Ordabayev for helpful discussions. We thank Haiqi Chen for helpful discussions regarding the mouse testis dataset. Components of our figures were created using BioRender. S.K. acknowledges funding through the National Science Foundation Graduate Research Fellowship under Grant No. DGE 2140743. F.C. acknowledges support from the Searle Scholars Award, the Burroughs Wellcome Fund CASI award, the Merkin Institute and the New York Stem Cell Foundation.

## Author Contributions

S.K., F.C., and M.B. designed the experiments. S.K. and L.D.A. designed and implemented the TissueMosaic model and software in consultation with M.B. S.K. and L.D.A. conducted data analysis. F.G. and S.F. contributed to the codebase. S.K., F.C., and M.B. wrote the paper with feedback from all other coauthors.

## Supplementary Figures

**Supplementary Figure 1.**
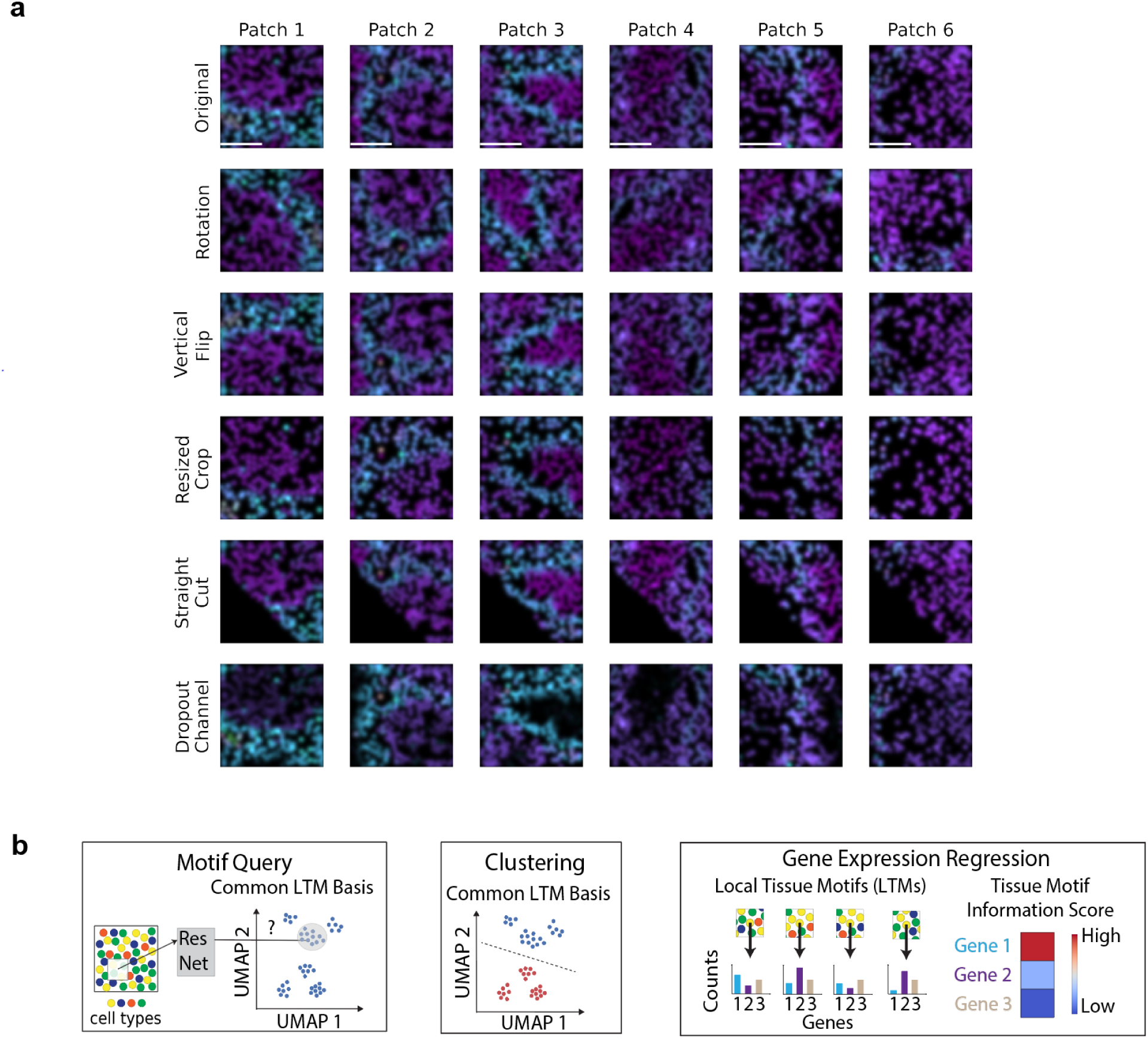
**a**. Examples of random augmentations applied during self-supervised training. Patches are randomly sampled from the Slide-seqV2 testis dataset. Spot dropout, horizontal flip, and multiplication by random global intensity are additional augmentations applied during training not shown here. During training, all augmentations are applied with certain probabilities. These probabilities and other specific augmentation parameters used during training can be found in Methods. Scale bar: 100 µm. Colors: Cell-type proportions transformed to RGB values based on *viridis* colormap. **b.** Self-supervised learning enables diverse downstream tasks such as motif query (finding similar motifs in the dataset to a certain query motif), clustering (grouping tissue motifs at a specific clustering resolution), and gene expression regression (identifying genes that have high mutual information with tissue structure in a cell-type specific manner).

**Supplementary Figure 2.**
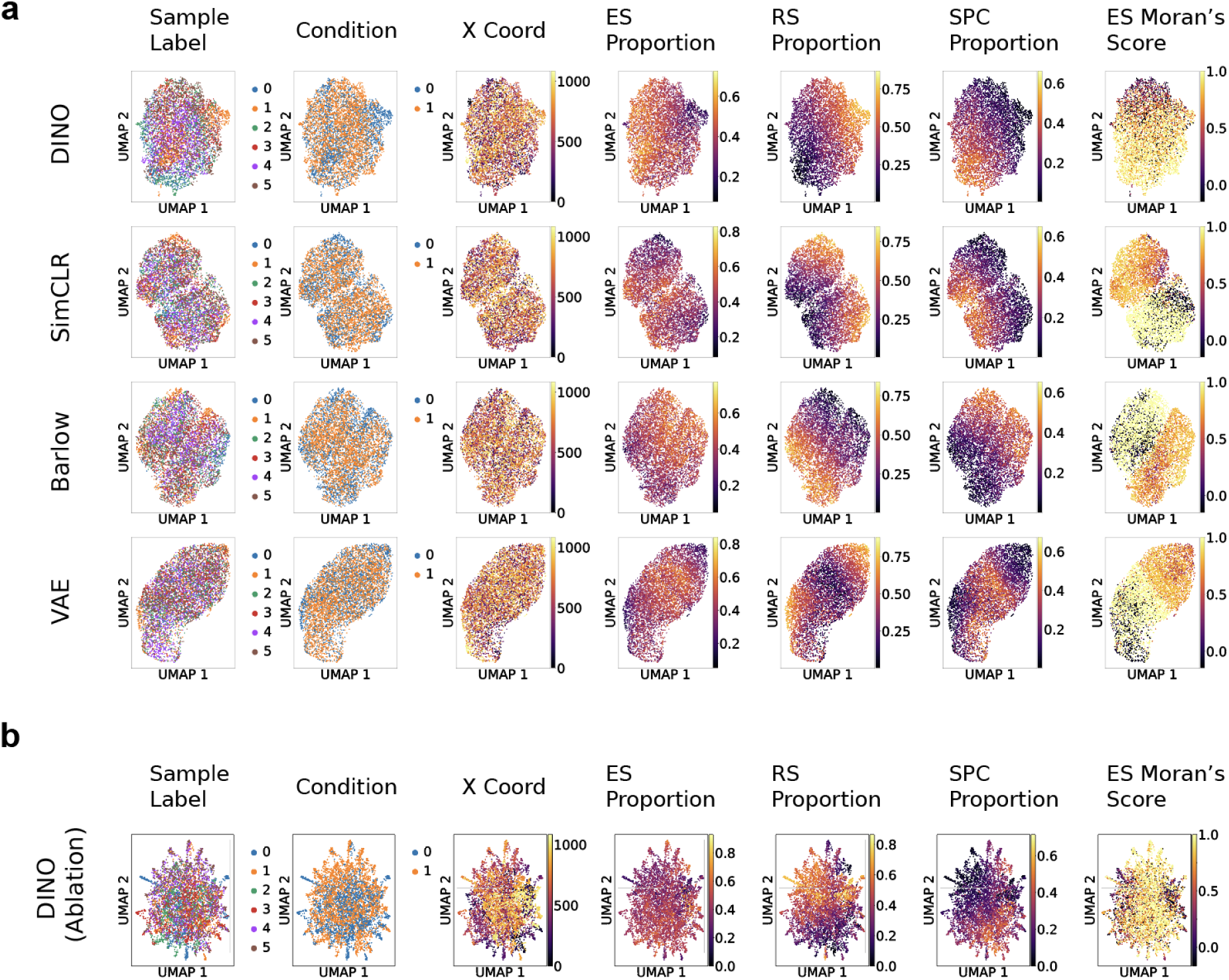
**a**. Uniform Manifold Approximation and Projections (UMAP) of patch representations for TissueMosaic trained with different self-supervised loss functions (DINO, SimCLR, Barlow) and variational autoencoder (VAE) baseline, colored by various features of interest (Colorbars represent column labels). Condition: 0 = wild-type, 1 = diabetes. “X Coord” refers to the x coordinate of the patch center. “ES, RS, SPC proportion” refer to the proportions of cells of each type respectively within the patch. “ES Moran’s Score” refers to spatial autocorrelation of ES cells computed by Moran’s I. **b.** UMAP of patch representations for TissueMosaic trained with modified DINO loss function, where local vs. global patch sampling strategy is ablated, and patches of single size are sampled as input to the two branches of the network.

**Supplementary Figure 3.**
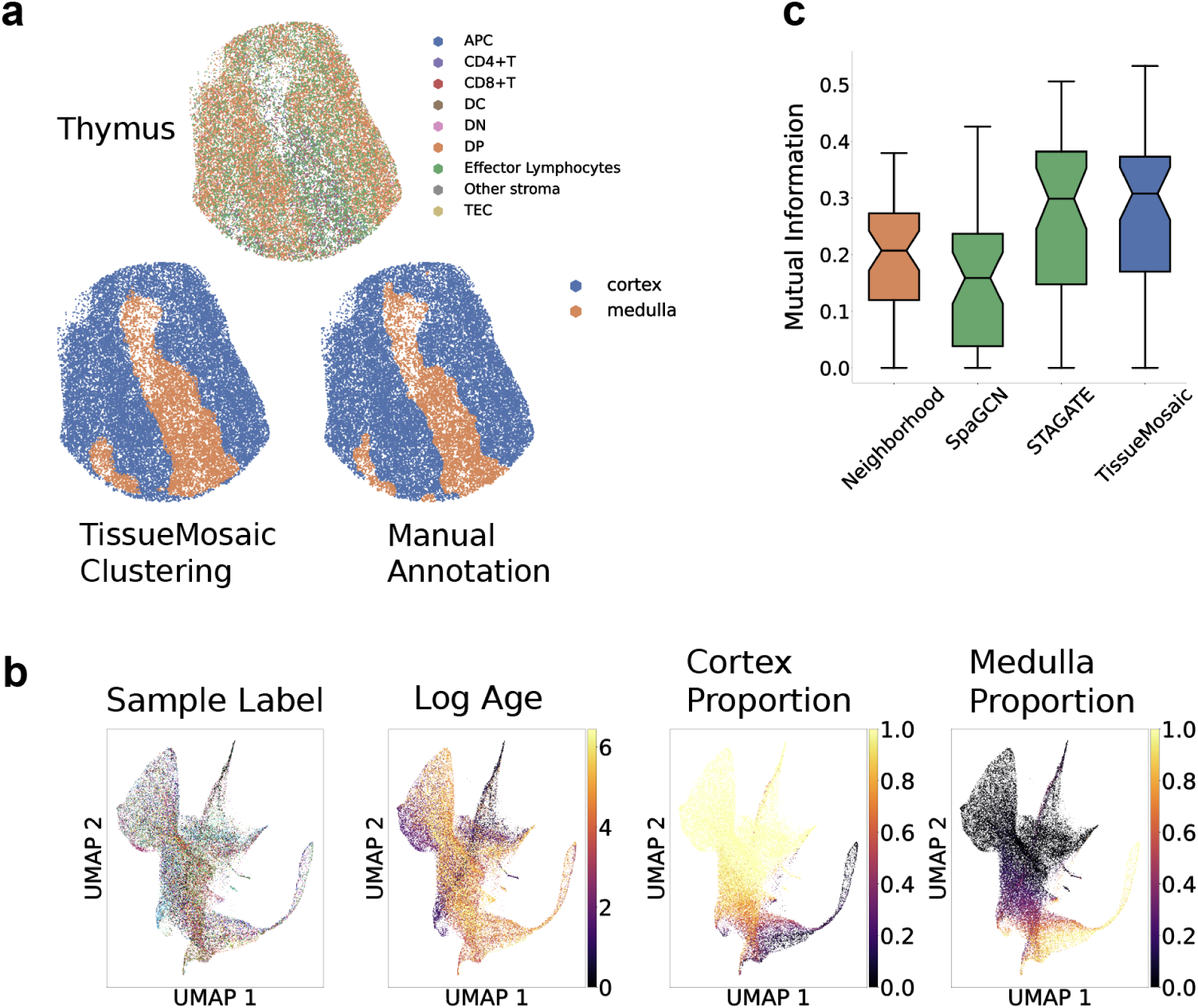
**a**. TissueMosaic clustering on a representative thymus sample. Cell-types, Top. Aggregated TissueMosaic clusters, bottom left. Manual annotation of same sample, bottom right. **b.** Uniform Manifold Approximation and Projection (UMAP) of patch representations for TissueMosaic (trained via DINO), Colorbars represent features of interest defined here:. Sample Label: 47 Slide-seqV2 samples, 1 color per sample. Log Age: Log2 of biological age in days. Cortex/Medulla proportion defined as fraction of beads in patch with manual annotation as cortex/medulla. **c.** Box plot of mutual information of clustering performance of existing spatial segmentation methods (SpaGCN, STAGATE), neighborhood composition baseline, and TissueMosaic across all thymus samples. Center = median, Box = IQR, Whiskers = farthest data point within 1.5IQR from box. Data beyond 1.5IQR are plotted individually. Notches indicate 95% confidence interval for median determined by bootstrapping. N=47 samples.

**Supplementary Figure 4.**
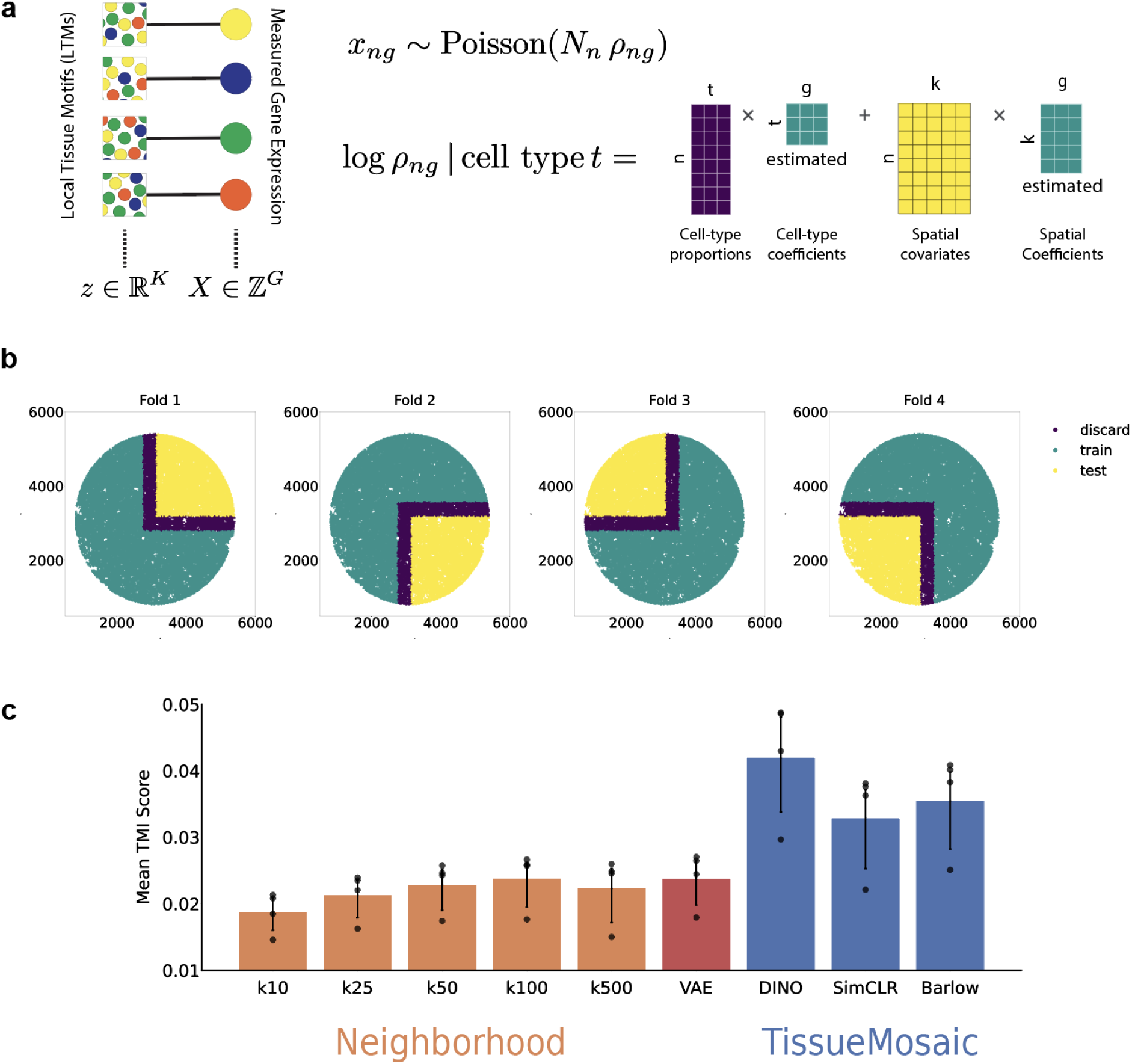
**a**. Poisson generalized linear model to predict measured gene expression (of dimensionality G) from cell-type proportions and spatial covariates (of dimensionality K, i.e. TissueMosaic representations). *N* is the number of measured UMIs; *n, g* index beads and genes respectively. **b.** Example of spatially partitioned train/test split strategy on a single Slide-seqV2 sample which produces 4 train/test folds with no spatial overlap between representations in the train or test splits. All spots are included in a test fold exactly once. **c.** Performance on gene regression task of neighborhood composition baseline across multiple neighborhood sizes, Variational Autoencoder (VAE) baseline, and TissueMosaic with various self-supervised loss functions (DINO, SimCLR, Barlow). Error bars indicate 95% confidence interval (determined by bootstrapping) of the mean Tissue Motif Information (TMI) score from spatially partitioned test folds (N=4, Methods).

**Supplementary Figure 5.**
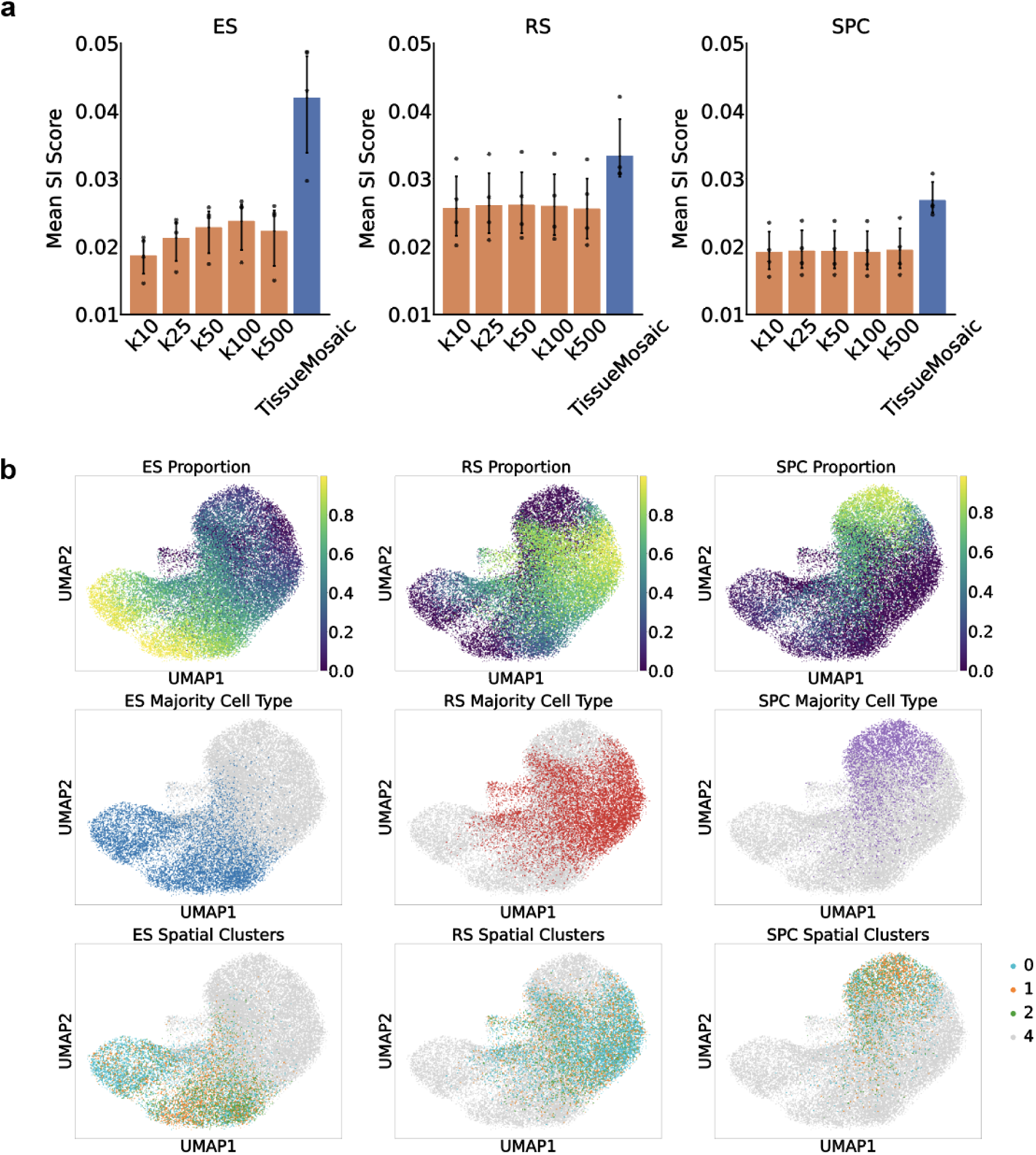
**a**. Performance on gene regression task across all wild-type samples for all three major cell types, using neighborhood composition baseline at multiple neighborhood sizes and TissueMosaic representations as spatial covariates. Error bars indicate 95% confidence interval (determined by bootstrapping) of the mean Tissue Motif Information (TMI) score from spatially partitioned test folds (N=4, Methods). **b.** UMAP of gene expression of each spot from representative testis sample, colored by cell-type proportions estimated by RCTD (top), majority cell-type labels (middle), and leiden clustering of TissueMosaic representations

**Supplementary Figure 6.**
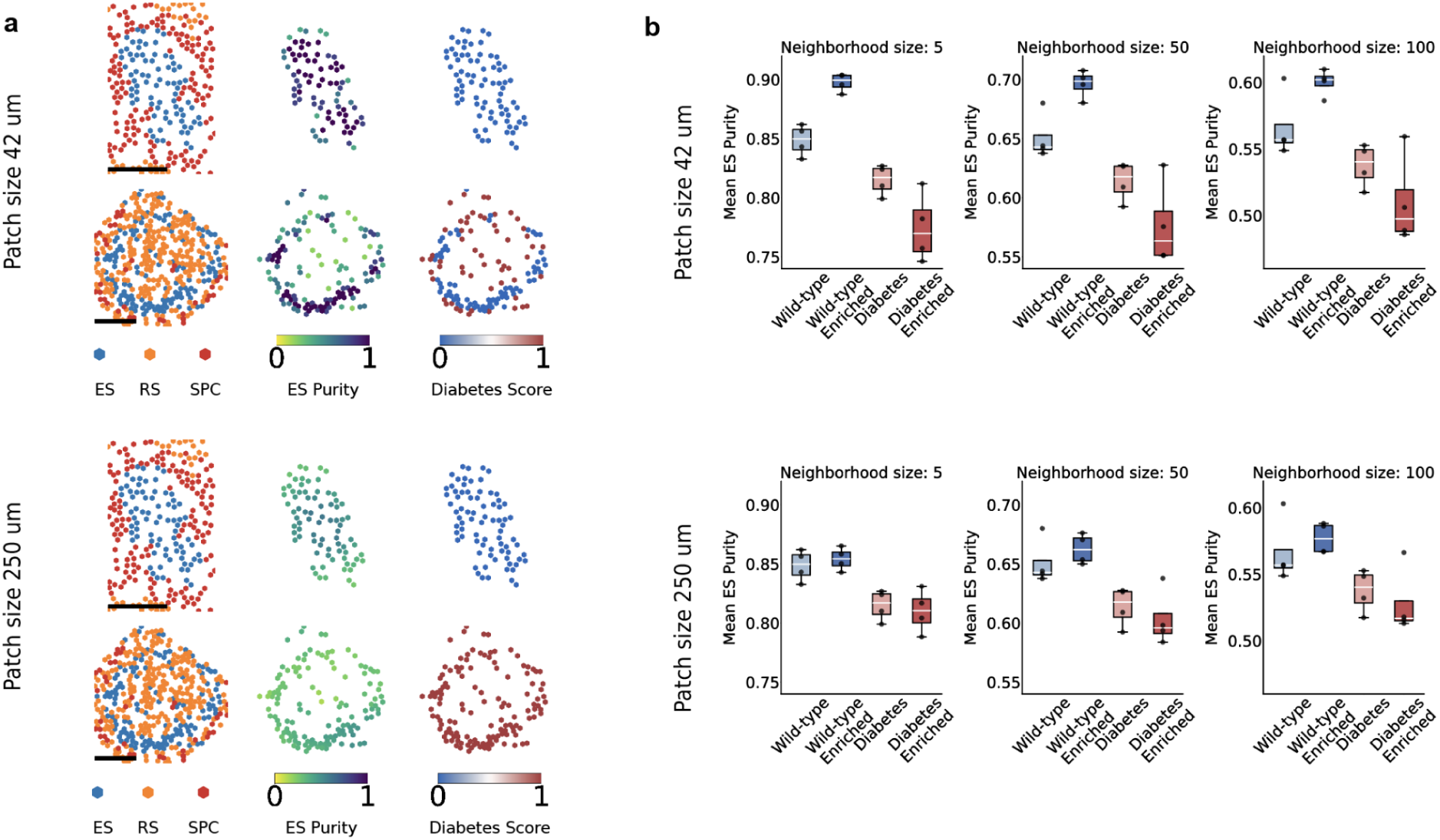
**a**. Example of conditional motif enrichment with varying patch size of model input on organized (above) and disorganized (below) tubules. Model input with patch size 42 um captures sub-tubular structure while model input with patch size 250 um captures super-tubular structure. Left, cell types. Middle, ES purity (defined as proportion of elongated spermatid cells in k Nearest neighborhood) calculated at neighborhood size 5 (top) and neighborhood size 100 (bottom). Right, TissueMosaic trained classifier prediction of ES status **b.** Box plots of mean ES purity within test fold (N = 4) before and after motif enrichment with varying patch size. Purity is calculated at multiple neighborhood sizes. Center = median, Box = IQR, Whiskers = farthest data point within 1.5IQR from box.

## Methods

### Model Architecture and Implementation Details

All self-supervised models are trained using a ResNet-34^28^ convolutional neural network (CNN) backbone with output layer of size 512, followed by a Multi-Layer Perceptron (MLP) projection head with two hidden layers of size 256 and 512 and output layer of size 512. The MLP projection head is only used during training and is discarded during inference. Adam optimizer is used with a minimum learning rate of 1.0E-5 and maximum learning rate of 5.0E-4, with 100 warm up epochs, 100 warm down epochs, and max epochs of 1000. In our experiments with the DINO SSL framework, initial student temperature of τ = 0.1 and initial teacher temperature of τ = 0.04 are used with momentum of m = 0.996 for both hyperparameters, and 2 global and 2 local crops are used. In our experiments with the Barlow Twins SSL framework, an off-diagonal penalty strength of λ = 0.005 is used. For the SimCLR SSL framework, a temperature τ = 0.1 is used.

### Augmentation Transformations

The self-supervised loss functions we implement rely on a set of random augmentations to train the underlying model. The model is encouraged to be invariant to the applied augmentations, and therefore the choice of augmentations is critical to learning useful representations. We choose a set of augmentations (**Fig S1A)** following those commonly used in computer vision along with ones based on specific properties of our spatial transcriptomics data. The augmentations are as follows:

*Dropout* – During preprocessing of spatial transcriptomics data, individual spots may be filtered out for quality control. To prevent sensitivity of the model to missing spots, we apply dropout to the cell-type image as a random augmentation. Dropout is applied with probability 1.0 and dropout rate is sampled from [0.1, 0.2, 0.3].

*Random Global Intensity* – The estimation of cell-type proportions that is used as input to the model is noisy; therefore, we multiply the cell-type image by a random global intensity to make the model less sensitive to this noise. This intensity is a user hyperparameter and for the results shown here is randomly sampled from (0.8, 1.2) and applied with probability 1.0.

*Random Rotation, Random Horizontal/Vertical Flip* – Random rotation, horizontal flip, and vertical flip which are commonly used augmentations in computer vision. Since rotating or flipping an image does not alter its intrinsic properties, our learned features should be invariant to these transformations. Horizontal and vertical flipping are applied with probability 0.5. Rotations are applied from –180 to 180 degrees with probability 1.0.

*Random Resized Crop* – Random resized crop is also a commonly used augmentation where a crop of the original image is made with a random area and random aspect ratio and then resized to a given size. This makes the model more robust to specific sizes and shapes of tissue motifs, within the scale of resizing. The random area of the crop is chosen from (0.8, 1.0) and the aspect ratio of crop is randomly sampled from (0.95, 1.05) before resizing to the desired patch size. For DINO only, for the local view, the random area of the crop is chosen from (0.5, 0.8) and aspect ratio is randomly sampled from (0.95, 1.05) before resizing to the desired local patch size.

*Random Straight Cut* – Slide-seqV2 data is collected on circular “pucks,” while the input patches to our convolutional neural network are square. Thus, patches sampled close to the edge of the puck will have portions of the patch that are empty. We therefore apply a random straight cut augmentation that occludes a specified fraction of the patch by zeroing all the data elements on one side of a random line passing through the patch. This augmentation encourages the model to learn representations that are invariant to the sampling location of the patch. Random straight cut is applied with probability 0.5, and the fraction of data that is occluded is randomly sampled from [0.1, 0.3].

*Drop Channel* – Different cell-types are often present in different abundances in spatial transcriptomics data, but the abundance of a cell type does not necessarily correlate with its biological importance. We therefore apply dropout of channels (cell-types) as a random augmentation. Channel dropout prevents the model from relying too heavily on any given cell-type (for example, the most abundant cell type). Channel dropout is applied with probability 0.2, with equal probability of dropout for all channels.

All augmentations are implemented in PyTorch to enable GPU acceleration or rely on the torchvision^29^ transforms library when applicable (specifically for Random Rotation, Horizontal/Vertical Flip, and Random Resized Crop).

### Model Training and Inference Workflows

The complete workflow during model training is as follows:

1. Populate tensor with input features (e.g. cell-type proportions) at corresponding coordinates from a spatial transcriptomics sample.
2. Randomly extract patches at 1.5x desired patch size.
3. Apply random dropout, random global intensity, and random rotation augmentations.
4. Center crop the patches to desired patch size.
5. Apply vertical flip, horizontal fip, random resized crop (for DINO only, this crop is resized to local size for student network), random straight cuts, and drop channel augmentations
6. Rasterize the tensor by convolving with a Gaussian kernel with σ randomly chosen from 1.0 or 1.5 pixels.

The complete workflow during inference is as follows:

1. Populate tensor with input features (e.g. cell-type proportions) at corresponding coordinates from a spatial transcriptomics sample.
2. Crop patch of desired patch size.
3. Rasterize the tensor by convolving with a Gaussian kernel with σ randomly chosen from 1.0 or 1.5 pixels.

### Evaluation Datasets, Data Preprocessing, and Dataset-specific Parameter Choices

We train and evaluate our model on two Slide-seqV2^5^ datasets of the mouse testis and thymus. The testis dataset consists of 3 wild-type and 3 diabetic (ob/ob) samples^23^. The thymus dataset consists of 47 samples across a time-course of mouse aging from day 0 to week 90, with 2-3 replicates per time point^13^.

For both datasets, cell-type proportions for each spot are determined using Robust Cell-Type Decomposition (RCTD)^9^, run in doublet-mode, which assigns 1-2 cell types per spot. Manual annotations of the cortex and medullary regions in the thymus dataset were determined by unsupervised clustering of the transcriptomes of each spot followed by iterative agglomeration of clusters to group clusters into cortex vs medullary regions and fine-tuned by visual comparison to H&E stains of serial sections^13^.

For mouse testis data, 1 pixel corresponds to 2.6 μm. Patches corresponding to global size 250 μm and local size 166 μm (only applicable to DINO) are extracted as input to the model for this dataset. For tissue motif enrichment on the mouse testis dataset, patches of global size 42 μm and local size 21 μm are used. For mouse thymus data, 1 pixel corresponds to 2.9 μm. Patches of global size 280 μm and local size 187 μm (only applicable to DINO) are extracted as input to the model for this dataset.

### Spatial Motif Query

To perform motif query on the testis, for each spot, we first obtain a TissueMosaic representation for a patch centered at that spot. The cosine similarity of the TissueMosaic representation for a query patch to all other patches in the dataset is computed and plotted as a transparency mask (**Fig 1B).** For benchmarking, STAGATE is run following the provided tutorial^17^ with default parameters and the produced representations are used for motif query. For quantification, the Pearson correlation coefficient (*r)* is computed between cosine neighborhood similarity and cosine representation similarity (similarity calculated between all query patches and all other patches in the dataset) using the numpy^30^ *corrcoef* function (**Fig 1C).**

### Spatial Clustering

To perform spatial clustering on the thymus, principal component analysis is first run on the TissueMosaic representations for a dataset. Uniform Manifold Approximation and Projection (UMAP)^31^ is run on the PCs that explain 90% of the variance of the TissueMosaic representations, with n_neighbors = 5000, min_dist = 0.5, n_components = 2, and metric = euclidean. Leiden^32^ clustering is performed at resolution of 0.2 with partition type RBC on the UMAP. Sub-clusters are aggregated into two spatial clusters for visualization purposes only (**Fig S3A)**. For benchmarking, SpaGCN and STAGATE are run following author provided tutorials with default parameters, except for similarly performing Louvain^32^ clustering at resolution of 0.2.

### Spatial Train/Test Split

We employ a spatial partitioning approach when we use the learned tissue motif representations for certain downstream tasks (**Fig S4B**). We divide each sample into 4 quadrants and for the Gene Regression and Motif Enrichment tasks, train 4 separate models, holding out one quadrant as a test quadrant each time. We discard any spots that are within a patch-sized border around the test quadrant from the train quadrant. This prevents any possibility of leakage from the train to the test set. All spots fall within a test quadrant exactly once. The regression or motif enrichment results for each test quadrant are then merged for downstream analysis.

### Gene expression regression

Gene expression regression is performed using the scikit-learn^33^ PoissonRegressor package. Input coefficients per spot index *n* are log number of UMIs 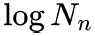, cell-type proportions 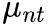, and optionally a set of spatial covariates 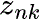:

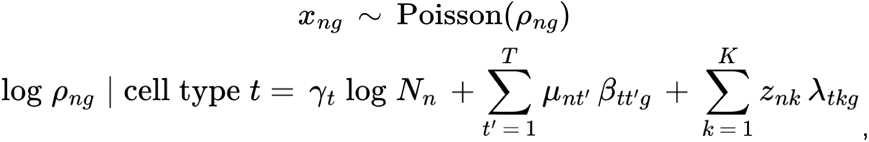

where *n* indexes spots, *g* indexes genes, *t* indexes cell-type, and *k* indexes spatial covariate dimension. Learnable coefficients are 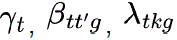.

For GLMs trained on neighborhood spatial covariates, 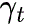 (log nUMI coefficients) are L2-regularized with a strength of α = 1.0e-6, 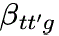 (cell-type proportion coefficients) with a strength of α = 1.0e-4, and 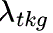 (spatial covariate coefficients) with a strength of α = 1.0e-2 or 5.0e-2. For GLMs trained on TissueMosaic or VAE spatial covariates, 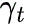 are L2-regularized with a strength of α = 1.0e-5, 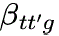 with a strength of α = 1.0e-3, and 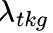 with a strength of α = 1.0e-1. These choices of L2 regularization strengths were determined by cross-validation.

In the testis dataset, spots with less than 100 UMIs are filtered out, and regression is performed for genes that are expressed in at least 10 percent of cells. In the thymus dataset, spots with less than 100 UMIs are filtered out, and regression is performed for genes that are expressed in at least 5 percent of cells. Any outliers in the TissueMosaic representations (element of feature vector > 2.0 in the learned representations of the testis data, element of feature vector > 3.0 in the learned representations of the thymus data) are set to 0 before performing regression.

### Evaluating Different SSL Frameworks

We train all three SSL frameworks, DINO^12^, Barlow Twins^11^, and SimCLR^2^ on a Slide-seqV2 dataset of mouse testis. We evaluate the performance based on the geometry of the learned latent space such as whether the representations are well mixed across samples and informative with respect to features of interest, such as cell type proportions within each patch (**Fig S2A)**. We also quantitatively evaluate each loss function using the gene regression task (**Fig S4C)**. We conduct evaluations on model checkpoints at intervals of every 100 epochs throughout the training process. For each loss function, we select the checkpoint that demonstrates the best performance. We find that through both evaluations, DINO leads to the most meaningful representations of our cell-type images.

### Motif Enrichment

We use the learned representations to train a classifier or regressor, employing a similar spatial partitioning strategy as used for the gene regression task. The classifier/regressor is trained using the sample level labels of patches from the train quadrants and then used to predict the status of patches from the test quadrants of each sample. These annotations are then assigned to the cell at the center of each test patch, to obtain an “enrichment” of the tissue into the desired phenotypes.

For the mouse testis dataset, motif enrichment is performed using a Ridge Classifier from the scikit-learn package on the model representations (of dimensionality 512). For the mouse thymus dataset, motif enrichment is performed using a Ridge Regressor. Ridge regularization strength is chosen via leave-one-out cross-validation from alphas of [1000, 2500, 5000], via the scikit-learn RidgeClassifierCV package and RidgeCV package respectively. Any outliers in the TissueMosaic representations (element of feature vector > 2.0 in the learned representations of the testis data, element of feature vector > 3.0 in the learned representations of the thymus data) are set to 0 before performing motif enrichment.

### Spatial Differential Expression

In the thymus dataset, differential expression analysis is run between young (motif enriched log age (days) < 5.0) and old (motif enriched log age (days) >= 5.0) antigen presenting cells using PyDESeq2^34^. Counts are pseudobulked based on motif enriched sample labels. PyDESeq2 performs a Wald test for differential expression and reported p-values are adjusted for multiple hypothesis correction across genes.^34^

In the testis dataset, to identify Δ*TMI* genes, two independent Poisson Generalized Linear Models are either trained on the elongated spermatid cells from wild-type samples vs diabetic samples (“Before motif enrichment”, **Fig 2g**) or the wild-type enriched (enrichment classification score < 0.5) vs diabetic (enrichment classification score > 0.5) elongated spermatid cells (“After motif enrichment”, **Fig 2g**). Δ*TMI* scores are obtained for genes that pass filtering criterion in both the wild-type and diabetic cells.

Gene set enrichment analysis (GSEA)^22^ is run with the M5 GO gene set^35^, including BP, CC, and MF components, with min_size=10, max_size=1000, permutation_num=10000.

